# Rapid-PFP: Accelerating Prefix-Free Parsing with GPU Parallelism

**DOI:** 10.64898/2026.04.28.721411

**Authors:** Eddie Ferro, Tyler Pencinger, Oded Green, Mahsa Lotfollahi, Christina Boucher

**Affiliations:** University of Florida, Gainesville, Florida, USA; NVIDIA Corporation, Santa Clara, California, USA

**Keywords:** GPUs, Prefix-free parsing, Succinct data structures, Data compression, Massively Parallel

## Abstract

Prefix-Free Parsing (PFP) is widely used in genomic data processing to construct compressed indexes on massive, highly repetitive datasets. However, existing CPU implementations are constrained by sequential bottlenecks, limiting their ability to scale to large-scale modern pangenomic collections. We introduce `RAPID-PFP`, a redesigned implementation of the `PFP` algorithm that takes advantage of the massive parallelism and high memory bandwidth of modern GPUs. `RAPID-PFP` parallelizes trigger-string detection, phrase parsing, dictionary construction, and parse generation through custom CUDA kernels and GPU-resident data structures built using cuDF, CuPy, and Numba-CUDA. The algorithm operates entirely within GPU memory, minimizes host interaction, and dynamically adapts to available VRAM, enabling efficient processing in a range of hardware configurations. Across E. coli and Human Pangenome (HPRC) datasets, `RAPID-PFP` produces identical output to established CPU pipelines while delivering an order-of-magnitude acceleration. On 3,682 E. coli assemblies, `RAPID-PFP` reduces runtime from 552 seconds to 17 seconds compared to `PFP-FL` (32.1 times) and from 1,078 seconds to 17 seconds compared to `PFP-ITL` (62.6 times). On the complete 46-sample HPRC dataset, `RAPID-PFP` achieves a 33.4 time speedup and successfully processes scales that `PFP-ITL` cannot handle. Performance improves with dataset size, reflecting that `PFP` maps naturally onto thousands of CUDA cores, yielding sublinear scaling relative to CPU implementations. `RAPID-PFP` demonstrates that foundational compressed-indexing algorithms can be re-engineered for accelerators, enabling scalable and practical preprocessing for large-scale genomic indexing workflows.

## 1. Introduction

Before the widespread adoption of high-throughput sequencing, sequencing and assembling a species’ reference genome was a resource-intensive endeavor. For example, the Human Genome Project, an international collaborative effort to produce the first human reference genome, cost approximately 2.7 billion (adjusted to 1991 USD) [19]. Today, technological advances have dramatically reduced sequencing costs, allowing large-scale projects to sequence thousands of individuals within a species, thus creating expansive pangenomic datasets. For example, the 1000 Genomes Project produced more than 6TB of assembled genomes [20], while the 100,000 Genomes Project aimed to sequence tens of thousands of genomes from individuals with rare diseases [21]. More recently, the Human Pangenome Reference Consortium (HPRC) has been working to create a comprehensive and diverse human pangenome reference by sequencing and assembling human genomes of diverse ancestry [11]. This proliferation of sequencing data extends beyond humans, with projects such as the 1001 Genomes Project [23] sequencing more than a thousand cultivars of *Arabidopsis thaliana*. The growth of these datasets challenges the use of a single reference genome and motivates a pangenomic approach, where multiple reference genomes collectively capture the genetic diversity of a species. However, this paradigm shift introduces computational challenges in the way reference genomes are selected and indexed, as the choice of reference has been shown to significantly impact alignment sensitivity [7, 8].

Prefix-Free Parsing (`PFP`) [5] was introduced in 2019 as a means to address these key computational challenges that arise in the analysis of large genomic datasets. `PFP` is a data-compression and preprocessing technique that transforms the input data into two components: a dictionary and a parse, which occupy significantly less space than the original data. This approach aligns with recent advances in dictionary- and grammar-based compression schemes that enable queries to be executed directly on the compressed representation. Unlike general-purpose compressors such as gzip, `PFP` preserves structural information required for efficient random access and downstream algorithmic operations. As a result, `PFP` can be used both to support direct queries on compressed genomic data [4] and to enable memory-efficient construction of a variety of compressed data structures [18, 6, 1, 3]. In practice, `PFP` has been used to enable scalable construction of pangenome indexes and succinct graph-based representations, and plays a key role in workflows where input can exceed hundreds of gigabases.

Although effective, current `PFP` implementations remain constrained by system-level limitations. The preprocessing pipeline contains sequential steps that restrict parallel speedup, and several stages require memory on the order of the input size, reducing the benefit of the compressed representation during construction. These factors limit performance in multicore architectures and hinder portability to accelerators and distributed-memory systems. Addressing these constraints is necessary for effective use of `PFP` in modern genomic pipelines, where data set sizes continue to increase and memory-efficient preprocessing is often the dominant cost.

In this paper, we present Rapid Prefix-Free Parsing (`RAPID-PFP`), a GPU-native implementation of `PFP` that re-engineers each stage for massive parallelism. The algorithm is designed to operate entirely within the GPU and requires minimal host resources. `RAPID-PFP` processes the input in chunks read directly into device buffers, launches a CUDA kernel to compute rolling hashes and mark trigger-string start positions, and parses all phrases in the chunk simultaneously by expanding boundary overlaps and constructing a cuDF string series. For each chunk, `RAPID-PFP` builds a temporary dictionary by grouping identical phrases and recording their occurrence indices; these temporary dictionaries are later merged, de-duplicated, and lexicographically sorted entirely on the GPU to finalize the global dictionary. Finally, `RAPID-PFP` constructs the parse by joining phrase occurrences with their position in the sorted dictionary and scattering the corresponding dictionary indices into the output parse array using a custom GPU kernel. This architecture avoids sequential bottlenecks, limits host-device transfers, and dynamically tunes chunk sizes to the available VRAM.

To evaluate the advantages of `RAPID-PFP`, we conducted an extensive experimental study comparing it with two widely used CPU implementations of `PFP: PFP-ITL`, released in 2018 [5], and `PFP-FL` [15, 16], a more recent and optimized implementation released in 2020. An overview of these methods is shown in Table 1. These experiments were performed on E. coli and Human Pangenome (HPRC) datasets of increasing size and complexity, enabling an assessment across realistic pangenomic workloads. Across all experiments, `RAPID-PFP` produced identical output to competing CPU methods, ensuring seamless compatibility with existing downstream tools. Our evaluation demonstrates that `RAPID-PFP` has an order-of-magnitude acceleration, achieving between 5.2 and 32.1 times acceleration over `PFP-FL` and between 22.3 and 62.6 times speedup over `PFP-ITL` on large E. coli datasets. On the largest benchmark (3,682 genomes), `RAPID-PFP` reduces runtime from 552 seconds (`PFP-FL`) and 1,078 seconds `(PFP-ITL`) to 17 just seconds, corresponding to a 96.7% and a 98.4% reduction in wall-clock time, respectively. On the Human Pangenome Research Consortium Year 1 release [11], `RAPID-PFP` is 33.4 times faster than `PFP-FL`, the only other method capable of completing on an input of this size. Most importantly, our results show that performance improvements grow with input size, reflecting how `RAPID-PFP` maps prefix-free parsing naturally to thousands of parallel CUDA cores. Unlike CPU-bound codes, whose runtime scales nearly linearly with input size, `RAPID-PFP` exhibits sublinear scaling, allowing datasets previously required hours of computation to be completed in less than a few minutes.

**Table 1.** Comparison of publicly available Prefix-Free Parsing (PFP) implementations, summarizing release year, code availability, and distinguishing algorithmic/engineering features (CPU vs. GPU execution, parallelism, and scalability).

| Method | Year | Repository | Notes / Characteristics |
| --- | --- | --- | --- |
| PFP-ITL | 2018 | https://gitlab.com/manzai/Big-BWT | First publicly available PFP implementation; part of Big-BWT pipeline; CPU-only; sequential parsing; known to encounter hash collisions on large inputs |
| PFP-FL | 2020 | https://github.com/marco-oliva/pfp | Optimized and modular C++ implementation; faster dictionary construction; supports multi-threading but remains CPU-bound; improved engineering over PFP-ITL. |
| RAPID-PFP | 2026 | https://github.com/EddieFerro/gpuPFP | Fully GPU-accelerated algorithm using CUDA kernels, CUDF, CuPy, and Numba-CUDA; massively parallelizes trigger-string detection, phrase parsing, dictionary building, and parse construction; operates entirely in GPU memory; adaptive batching enables use on GPUs with varying VRAM; produces identical output to CPU versions. |

In summary, our main contributions are as follows.

- **First end to end GPU implementation of PFP, to our knowledge**. We present `RAPID-PFP`, the first GPU native realization of the complete PFP pipeline, showing that a compression aware preprocessing algorithm originally engineered for CPU workflows can be redesigned to exploit thousands of CUDA cores and high memory bandwidth.
- **Full pipeline parallelization using GPU resident data structures**. `RAPID-PFP` parallelizes every stage of `PFP`, including trigger string detection, phrase materialization, dictionary construction, and parse generation, using custom CUDA kernels and GPU libraries such as cuDF and CuPy, while producing bit identical output to established CPU implementations for seamless downstream compatibility.
- **VRAM aware design and order of magnitude empirical gains at pangenome scale**. `RAPID-PFP` is engineered around GPU memory constraints by streaming inputs in adaptively sized chunks, minimizing host device transfers, and requiring minimal host resources. Across *E. coli* and HPRC workloads, this design yields order of magnitude runtime reductions relative to state of the art CPU implementations, with speedups that increase with dataset size.

All of our code is publicly available on GitHub at https://github.com/EddieFerro/gpuPFP.

## 2. Preliminaries

In this section, we provide a comprehensive overview of Prefix-Free Parsing (`PFP`), a preprocessing algorithm. Before presenting the details of `PFP`, we outline the foundational definitions necessary to understand its mechanics.

A string *T* is defined as a finite sequence of symbols, *T* = *T* [1..*n*] = *T* [1] ⋯ *T* [*n*], over an alphabet Σ = {*c*_1_, …, *c*_*σ*_}, where the symbols can be ordered lexicographically unambiguously. The substring of *T* from position *i* to *j* is represented as *T* [*i*..*j*], where *i ≤ j*. Given two strings *T*_1_ and *T*_2_, their concatenation is defined as *T*_1_ + *T*_2_, which appends all symbols of *T*_2_ to *T*_1_. We define a string of length *k* composed of a single character *c* as *T* = *c*^*k*^.

For a string *T* of length *n, T* [1..*i*] is called the *i*-th prefix of *T*, and *T* [*i*..*n*] is called the *i*-th suffix. A prefix *T* [1..*i*]is a *proper prefix* if 1 *≤ i* < *n*, and a suffix *T* [*i*..*n*] is a *proper suffix* if 1 < *i ≤ n*. The lexicographic order between two strings *T*_1_ and *T*_2_ is denoted by ≺. Specifically, *T*_2_ ≺ *T*_1_ if *T*_2_ is a proper prefix of *T*_1_, or there exists an index 1 *≤ i ≤* min(|*T*_1_|, |*T*_2_|) such that *T*_2_[1..*i* − 1] = *T*_1_[1..*i* − 1] and *T*_2_ [*i*] < *T*_1_ [*i*].

### 2.1. Rank

We also use the term *rank* to refer to a string’s position in a lexicographically sorted list. Formally, given a set (or list) of distinct strings *S* sorted in increasing lexicographic order, the rank of a string *x ∈ S*, denoted rank(*x*), is the unique index *r* such that *x* is the *r*-th element of the sorted order.

For example, let *S* = {GCCA, ACCA, TCCA}. Sorting lexicographically gives *S* = ⟨ACCA, GCCA, TCCA⟩ . Thus rank(`ACCA`) = 1, rank(`GCCA`) = 2, and rank(`TCCA`) = 3.

### 2.2. GPU Architecture

A typical GPU architecture is designed to support massive parallelism through thousands of lightweight threads organized into a hierarchy of processing units. At the core of this structure are Streaming Multiprocessors (SMs), each capable of executing many threads concurrently using the Single-Instruction, Multiple-Thread (SIMT) model. Threads are grouped into warps—typically 32 threads per warp—which execute instructions in lockstep. Each SM contains its own register file, shared memory, and L1 cache, which allow low-latency communication and data sharing between threads within a block. GPUs also include global memory and L2 cache shared across all SMs, allowing broader data exchange and persistent memory access. Efficient memory access patterns and hardware-managed thread scheduling allow GPUs to hide latency and sustain high throughput for compute- and memory-intensive tasks. Communication within a thread block is facilitated via shared memory, while communication across blocks relies on global memory, which is slower but accessible to all threads.

GPU kernels, or programs to be executed by a GPU in parallel, are launched on the device as a one-dimensional grid of blocks, where users define the number of blocks and the number of threads per block (up to a maximum of 2048 threads per block). During execution, each block in the grid is assigned to a SM which executes the kernel in every thread in the block by scheduling the warps in each block. Each SM executes the warps in a round-robin fashion and switches between the different blocks assigned to it to maintain high-utilization and avoid any latency. The kernel is complete once every thread has executed its instructions. In practice, it is important to choose enough blocks so that every SM is busy and to choose enough threads so that the memory latency of warps can be hidden by switching, while avoiding so many threads that it exhausts shared block resources. Figure 1 shows the typical structure of a GPU 1. Note that each block typically contains more than one warp. In our paper, we ran all experiments on a single NVIDIA B200 GPU, which is based on the Blackwell architecture. It comprises 144 SMs, each capable of handling up to 2048 concurrent threads. Each SM in the B200 includes 128 FP32 CUDA cores and 4 fifth-generation Tensor Cores, in addition to local shared memory and L1 cache. Threads communicate within blocks using fast shared memory, while interblock communication occurs through a high-speed 60MB L2 cache and HBM3e global memory, which offers up to 8.0 TB/s bandwidth and 192GB of memory.

**Figure 1:**
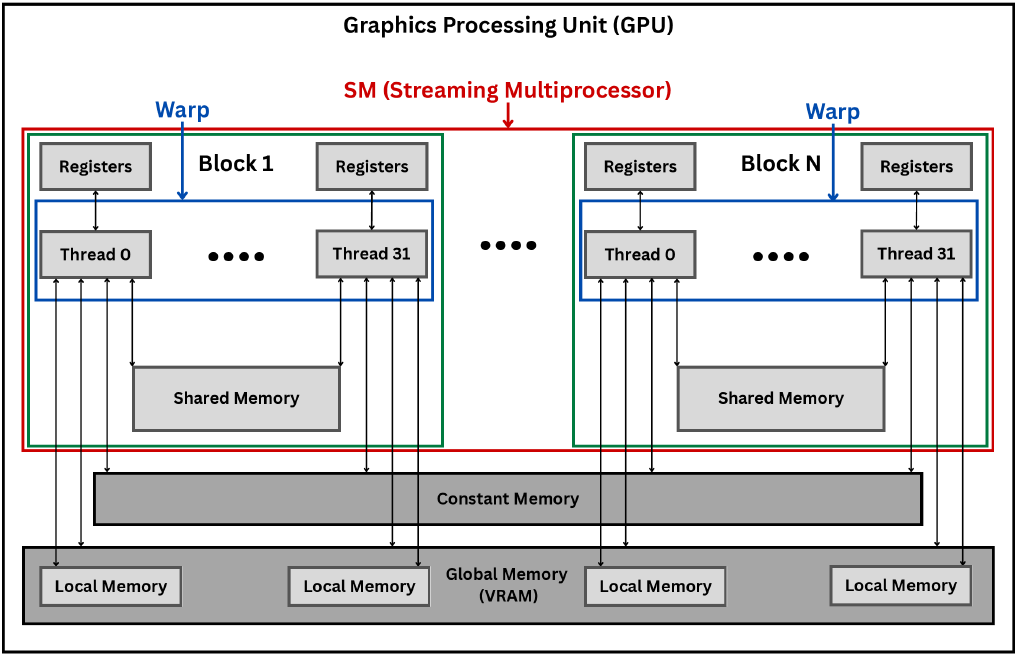
GPU Memory and Execution Hierarchy.

### 2.3. CUDA Data Types and Functions

We use a number of CUDA data types and functions that we briefly describe here. First, we use the cuDF DataFrame library [17]. This library is centered around Series, which are one-dimensional labeled arrays of data, and DataFrames, which are two-dimensional collections of Series with aligned rows and labeled columns. Within the context of cuDF, a Series is stored as a contiguous GPU array in a buffer, and a DataFrame, being essentially a collection of Series, has all of its columns stored as a contiguous GPU array in their own buffers. We note that data types in cuDF are strongly typed, and each Series can store only one data type, but a DataFrame can store a different data type in each of its columns.

cuDF operations are performed in parallel using the GPU’s architecture, enabling rapid execution of common tasks such as filtering, sorting, joining, grouping, aggregating, and applying user-defined functions. cuDF minimizes host-device data transfer by keeping the data resident in GPU memory, thereby reducing latency, and improving overall throughput. In particular, cuDF offers functions such as `concatenate, drop_duplicates, and sort_values` that can perform operations on an entire DataFrame or Series at the same time through the GPU.

We also use the CuPy GPU Array Library [14] which is analogous to the NumPy and SciPy [9, 22] libraries available for Python. CuPy allows arrays to be allocated in GPU memory and perform element-wise and linear algebra operations on the arrays. These operations are executed by the library through optimized CUDA kernels that accelerate the operations with parallelization on the GPU. CuPy also serves as the foundation for the *cuDF DataFrame library*. This library is primarily used when the full functionality of cuDF is not necessary and CuPy provides the necessary operation itself. One specific operation is the where operation that identifies all indices in an array where a user-specified condition is specified. We use the *Numba-Cuda* just-in-time (JIT) compiler [13] to write Python kernels that can be executed on the GPU. Additionally, we use the *RAPIDS Memory Manager (RMM)* library [17] to allow the libraries mentioned above to share a unified pool of memory that is managed by an RMM allocator.

### 2.4. Prefix-Free Sets

To introduce the key concept underpinning `PFP`, we define *prefix-free sets* that play a central role in the algorithm’s correctness.

#### Definition 1

(Prefix-Free Set). A set of strings 𝒮 is *prefix-free* if no string in 𝒮 is a proper prefix of another string in 𝒮.

For example, the set 𝒮 = {ACCA, GCCA, TCCA} is prefix-free. This property will be used throughout the paper to ensure that phrase boundaries and their corresponding suffixes can be handled unambiguously in `PFP`.

### 2.5. Prefix-Free Parsing

`PFP` is a lossless preprocessing algorithm that can parse large strings into a compressed representation. `PFP` takes as input a string *T* of length *n*, and two integers greater than 1, which we denote as *w* and *p*. It parses *T* into unique overlapping phrases, which are stored in a dictionary in their lexicographic ordering. A parse is then constructed that contains the order in which the phrases appear in the original input, where each phrase is represented by their position in the dictionary. The input can then be reconstructed by iterating through the parse and concatenating the phrases in the order indicated by the parse, while accounting for the overlap between phrases.

We denote the dictionary by `D` and the parse by `P`, and note that `D` is referred to as a dictionary, but in practice it is stored on disk as a string with all phrases concatenated together with a special symbol separating them. As the name `PFP` suggests, this output has the property that none of the suffixes of length greater than *w* of the phrases in `D` is a prefix of any other. We formalize this property through the following lemma. We refer to `PFP` of *T* as `PFP`(*T*), consisting of the dictionary `D` and the parse `P`.

#### Lemma 1.

*[5] If we are given a string T and* `PFP`(*T*) *then the set 𝒮 of distinct proper phrase suffixes of length at least w of the phrases in* `D` *is a prefix-free set*.

The first step in `PFP` is appending *w* copies to *T* of a special symbol that is lexicographically smaller than any character in Σ which we will denote as $. For simplicity, we consider the string *T*′ = $^*w*^*T* $^*w*^. Next, we characterize the set of trigger strings `E`, which will be used to define the parse of *T* . Given a parameter *p*, we construct the set of trigger strings by computing a rolling hash (e.g. Karp-Rabin hash [10]), *H*_*p*_(*t*), of all substrings of length *w* in *T*′. Then, the set `E` is defined as the set of substrings *t* = *T*′[*s*..*s*+*w*−1], where *H*_*p*_(*t*) ≡ 0 or *t* = $^*w*^. This set of trigger strings is then used to parse *T*′ by iterating through every trigger string starting position and taking a substring starting at one of these positions and ending at the end of the next trigger string. If we denote *P* as the list of starting positions of the trigger string, then every substring, except the last, is defined as *T* [*P* [*i*]..*P* [*i* + 1] + *w*]. This process turns the input into a set of phrases that have an overlap of *w* with at least one other phrase. The dictionary is the set of all unique phrases in their lexicographic order that were identified in this manner. The parse then is an integer list which describes the order in which all the phrases appear in the input by referencing each position of each phrase in the sorted dictionary.

We now walk through these steps on the toy example

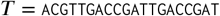

with *w* = 2. Since *w* = 2, we first pad the original string *T* on both sides with $$ to obtain

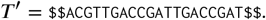

Next, we slide a window of length *w* = 2 across *T*′ and compute a rolling hash for each 2-mer. A 2-mer *t* = *T*′[*s*..*s* +*w*−1] is declared a *trigger string* if either (i) *t* = $$ (the boundary trigger), or (ii) its hash satisfies *H*_*p*_(*t*) ≡ 0 (mod *p*). For example, suppose we end up with the following set of trigger strings:

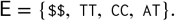

The dictionary *D* is the set of all unique phrases, sorted lexicographically. In this example,

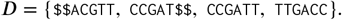

Finally, the parse is the sequence of dictionary ranks that reproduces the phrase order in the input. Using the phrase list above, we obtain

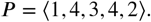

To reconstruct *T*′ from (*D, P*), we concatenate the phrases in parse order while removing the *w*-character overlap between consecutive phrases. Since each phrase ends with the next trigger string, this overlap removal is well-defined and yields exactly *T*′. In summary, `PFP` transforms *T*′ into a compact pair (*D, P*) that captures repeated structure via phrase reuse, and this representation supports efficient construction of downstream indexes.

### 2.6. PFP CPU Implementation

The current implementation of PFP on the CPU is inherently sequential. It starts by initializing a hash map to represent the dictionary. Then, it processes a single sequence at a time. For each sequence, it iterates over every character and adds it to the phrase that is currently being built. It also updates the Karp-Rabin hash to be over the last *w* characters of the phrase that is being built. When the Karp-Rabin hash of the current window meets the criteria for a trigger string, the current phrase is put into the dictionary hash map if it is not already there. Then, the hash used to store this phrase in the hash map is immediately written out to a temporary parse file. The execution continues with the construction of the next phrase which already starts with the last *w* of the just-processed phrase. Once all sequences have been handled in this sequential manner, the phrases and their hashes are taken from the dictionary hash map, and the phrases are sorted lexicographically. Finally, the parse is streamed in from the temporary file, a lookup is performed to relate the incoming hash to the position of the corresponding phrase-hash pair in the sorted dictionary, and then the position index is streamed out to the final parse file. Finally, the dictionary is written to disk without any of the hash values, just the phrases.

## 3. RAPID-PFP Algorithm

### 3.1. Overview

The `RAPID-PFP` algorithm is broken up into three main components: the PFP step, the dictionary finalization step, and the parse construction step. The PFP step applies each step of the PFP algorithm in a massively parallel fashion to chunks of the input and then outputs an intermediate result for processing in the later steps. For example, in the CPU implementation trigger strings and phrases are found at the same time by iterating over the input, but in `RAPID-PFP` all trigger string positions in a chunk are found at once with a kernel and all the phrases in the chunks are parsed at once with another kernel. Both the dictionary finalization and parse construction steps use the intermediate results from all of the chunks processed by the first step to produce the final dictionary and parse output.

### 3.2. Data Input step

`RAPID-PFP` uses the KvikIO library to read chunks of data from the input file directly to a GPU buffer with minimal Host-side overhead, which allows for efficient data input. By default, chunks equal to 10% of the available VRAM on the device are read one at a time and processed. This allows enough space in device memory for multiple additional data structures, usually equal to the size of the chunk, that are necessary in the PFP step to exist in GPU memory at the same time. This value can be changed by the user. Additionally, *w* copies of a special character smaller than any other character that occurs in the input as required by the PFP algorithm are prepended and appended to the chunk.

### 3.3. PFP step

#### 3.3.1. Trigger String Finding

After an input chunk has been read, the first step of the `RAPID-PFP` algorithm is to identify all the trigger string positions. This is done using a custom Numba CUDA kernel, which uses the Numba JIT compiler to transform Python code into an LLVM intermediate representation and then into native device code. The kernel, `trigger_string_finder`, takes in the input chunk buffer, the user defined values for *w* and *p*, a prime number (smaller than 2^32^ − 1), and a results buffer of the same size as the input chunk buffer. During the kernel execution, every thread uses its absolute position within the grid to identify which position it corresponds to in the input. Then, it computes a polynomial hash, using the given prime, of a window of size *w* starting from its position in the input. For example, the thread with absolute position 1 (thread 1 in block 1) corresponds to position 0 of the input chunk and computes the hash for the window *T* [0..*w*]. The remainder of the hash value when divided by *p* (hash mod *p*) is used to determine whether the window meets the trigger string condition. If the remainder is equal to zero, or the window is the first or last window of the input chunk, then the same position as the start of this window in the input is marked with a 1 in the result buffer. For example, the thread with absolute position 1 will output a 1 in position 1 of the output buffer. Therefore, the result buffer produced by this kernel encodes, for each position in the input chunk, whether that position is the beginning of a trigger string. Note that this encodes essentially what a bit array would encode but does not offer any of the functionalities of a bit array and takes more space. The kernel is executed with 128 threads per block and enough blocks so that there is a thread for each window in the input, which can be determined at runtime. The pseudocode for this kernel can be found in Algorithm 1.

##### Algorithm 1

Trigger String Finder CUDA Kernel

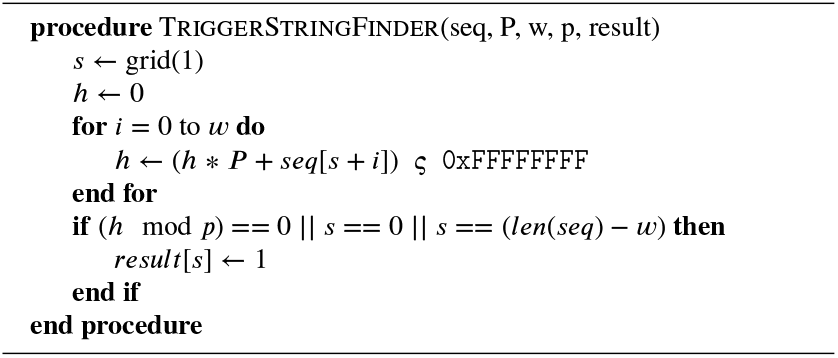

#### 3.3.2. Phrase Parsing

Using the output buffer from this kernel, we can efficiently parse every single phrase in the input chunk. To do this, we first use the CuPy *where* function to find all the indices in this output buffer that contain a value of 1. These indices represent the start position of every trigger string in the input chunk and since every phrase starts with a trigger string, these indices (except the last) also mark the start of every phrase in the input chunk. Additionally, every phrase must share a trigger string with the phrase that follows it, so the end position of every phrase can be found by adding *w* to the start position of its following phrase. For example, lets say *w* = 2 and the first three indices are [0, 12, 36], then the first phrase would be from *T* [0..14] and the second phrase would be from *T* [12..38].

Using these indices would allow us to parse the input chunk directly on the GPU by constructing a cuDF Series where each phrase is a row in the Series. This is because internally a string Series is represented on the GPU by a char buffer of all the strings concatenated together and an integer buffer whose values denote the separation between these strings in the char buffer. However, every offset value in the integer buffer (except the first and last) marks the end of one string and start of another. This does not allow us to define any overlap between subsequent phrases in the input chunk when parsing with a cuDF series construction. The solution is to expand the sequence at every trigger string (except the first and last) by duplicating the trigger string and shifting the start indices accordingly. Let’s say we have the input chunk *T* = $$*ACGT ACGT ACCT AGACA*$$ with indices [0, 3, 7, 12, 20] and *w* = 2, then the expanded sequence and updated indices would be *T* = $$*ACGCGT ACGCGT ACCT CT AGACA*$$ and [0, 5, 11, 18, 26]. Note that the sequence is expanded by (*k* − 1) * *w*, where *k* is the number of phrases, and each index (except the last) is shifted by *i* * *w*, where *i* is the offsets position in the buffer.

In order to compute the shifted indices, we use a vectorized subtraction of subsequent start indices given by the CuPy *where* function and add *w* to the resulting values with a vectorized addition to get the length of each phrase in the input chunk. Then, we utilize the CuPy *cumsum* function to compute the exclusive prefix sum of the lengths of the phrases to get our final shifted indices. Formally, the shifted indices, *I*, are defined as *I*[0] = 0 and *I*[*k* + 1] = 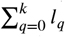, where *l*_*q*_ is the length of the *q*-th phrase in the input chunk. With the shift, every index now represents both the end and start of the phrase. Then, in order to create the expanded sequence, we use a custom Numba CUDA kernel, `expand_sequence`. This kernel takes in the original input chunk buffer, the buffer containing the original indices, the size of the indices buffer, the value *w*, and the buffer where the expanded sequence will be stored. It spawns a thread for every single position in the expanded sequence buffer and, at a high level, every thread will determine which position in in the original sequence, *i* it’s position in the expanded sequence, *j*, corresponds to and copy that value over to the expanded sequence buffer. Each thread determines this by invoking a device-side custom binary search kernel, _`upper_bound_duplications`, that determines the number of duplicated trigger strings have occurred at or before its position in the expanded sequence with a binary search over the buffer of the original indices. The pseudocode can be found in Algorithm 2.

##### Algorithm 2

Upper Bound Duplicates CUDA Kernel

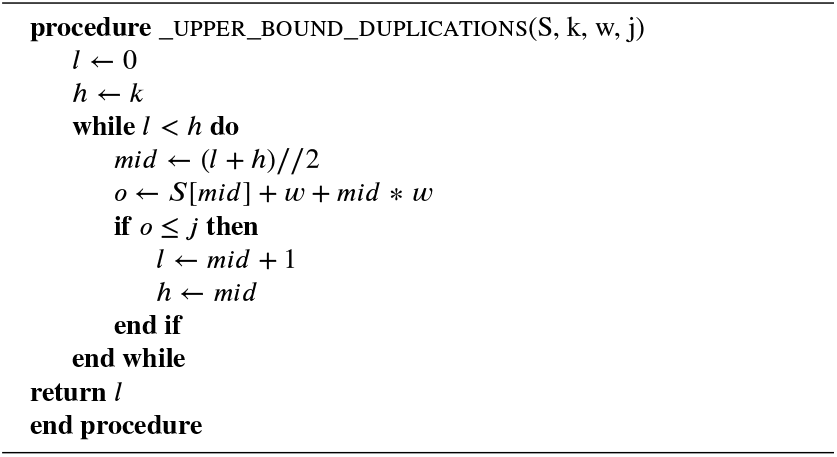

Once each thread determines the number of duplications that have occurred before or at its position, *u*, it can determine which value it is responsible for carrying over. In the case that no duplications have occurred, then the thread directly copies over the value from the same position in the input (*out*[*j*] = *in*[*i*], where *j* == *i*). If duplications have occurred, then each thread determines the start position in the expanded sequence of the last trigger string to be duplicated before the thread’s position in the output. Let’s assume that *t* is the index of this trigger string in the buffer of original indices, *S*, then the start position of the duplication of this trigger string, *x*, in the expanded sequence is given by *x* = *S*[*t*] +*w*+*t* * *w*. The thread then distinguishes whether its position falls within a duplicated trigger string (*j* ∈ [*x, x* + *w*)) or if it is part of the shifted original sequence. If *j ∈* [*x, x*+*w*), then its corresponding position in the original input is *i* = *S*[*t*] + (*j* − *x*). Otherwise, its corresponding position is *i* = *j* − *u* * *w*. This kernel is executed with 128 threads and enough blocks so that there is a thread for each position in the expanded sequence. The pseudocode can be found in Algorithm 3. Finally, a cuDF Series is defined using the expanded sequence and the shifted indices so that the input chunk is parsed into all of its phrases.

##### Algorithm 3

Expand Sequence CUDA Kernel

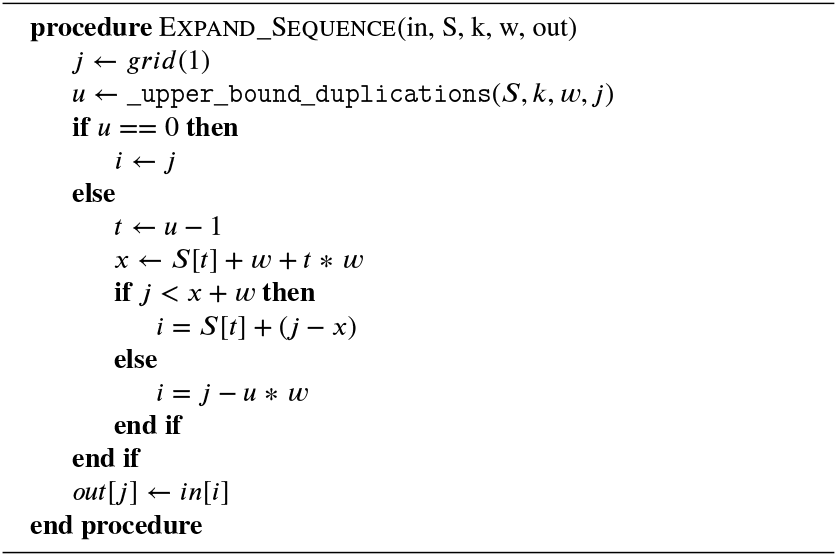

#### 3.3.3. Temporary Dictionary Construction

Once each input chunk is parsed into phrases as a cuDF Series a temporary dictionary is built using this Series and written out to a temporary parquet file. To construct the temporary dictionary, we place the Series into a Dataframe as the phrases column and use the `reset_index` function to create a numeric column called index so that each phrase in the phrases column has a corresponding index value in the index column. The maximum value in the index column is found using the `max` function and used to update a cumulative offset value, and the index column is incremented by the current cumulative offset value to keep indices globally unique across chunks. Then, we use a `groupby` operation on the phrases column alongside a `collection aggregation` on the index column to remove duplicate phrases and collect a list of all associated indices with each phrase. This is written to a temporary parquet file with the `to_parquet` function.

### 3.4. Dictionary Finalization Step

In the final dictionary step, we consolidate all the temporary dictionaries that were created for each input chunk. This is done by using the cuDF `read_parquet` function to read in the phrases column from the temporary dictionary Dataframes that were written to parquet files in the previous step. As each parquet file is read in, it is concatenated to the final dictionary Dataframe that is being constructed. Then, duplicate phrases that may have occurred across input chunks are removed at this time using the cuDF `drop_duplicates` function. Once all the temporary parquet files have been read in and all duplicates dropped, the dictionary is sorted with the `sort_values` function. Finally, the dictionary is written out in the expected format of `PFP` using the `to_csv` function. There is also an option to output the dictionary in a parquet format, which takes less space on disk than the expected dictionary format and also takes less time to output. Note that the Dataframe containing the final dictionary is maintained in GPU memory for the next step.

### 3.5. Parse Construction Step

To construct the parse we need to consolidate all the indices for each phrase and ensure that each phrase’s rank in the sorted dictionary is output to its indices. First, we allocate the buffer for the parse with a total length of the cumulative offset value that was updated at every temporary dictionary construction, which is equivalent to the total number of phrases in the input file after applying `PFP`. Next, we create a numeric column called rank in the dictionary Dataframe to relate each phrase to its rank in the sorted dictionary. Then, we read in the phrases and index columns of each temporary parquet file into a Dataframe and perform a `merge` between the temporary parquet Dataframe and the sorted dictionary Dataframe. This creates a relationship between each phrase’s indices which were read in from the parquet file and its rank which is part of the dictionary.

Using this information, we can fill out the parse buffer by placing each phrase’s rank at all of it’s indices. To do this efficiently, we utilize a custom Numba CUDA kernel called `scatter_row_index`. It takes as input: a positions buffer, an offsets buffer, a ranks buffer, and the parse buffer. The positions buffer is a contiguous buffer of all the positions of all the phrases in the temporary parquet Dataframe, which is obtained by decomposing the lists of positions of each phrase. The offsets buffer indicates which positions in the flat positions array belong to which phrase and is obtained by computing an exclusive prefix-sum over the lengths of each phrase’s positions list. The ranks buffer simply relates each phrase to its rank. The kernel spawns a thread for every phrase in the temporary parquet Dataframe and each thread is responsible for outputting it’s corresponding phrase’s rank to the same phrase’s positions. This kernel’s pseudocode can be seen in Algorithm 4. After every temporary parquet file has been processed and the parse buffer has been fully populated, it is written out in the expected format of `PFP` using the KvikIO library.

#### Algorithm 4

Scatter Row Index CUDA Kernel

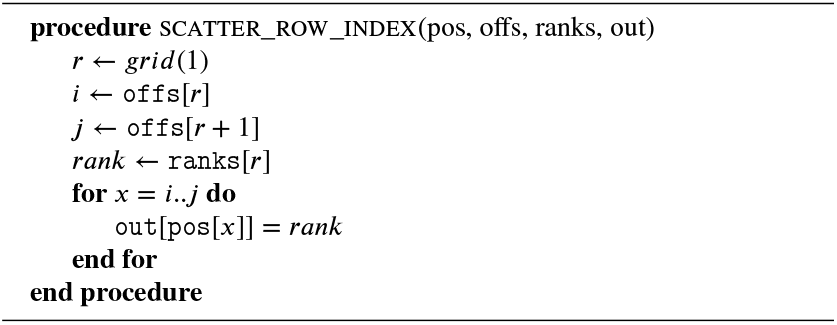

## 4. Results

### 4.1. Datasets

Our experimental evaluation used two datasets. The first set of datasets consisted of concatenated *E. coli* assemblies from NCBI [2], with the number of assemblies increasing for subsequent datasets. In particular, we concatenated subsets of these assemblies to create files of 500, 1,000, 2,000, and 3,682 concatenated assemblies. We will refer to these datasets as `ecoli.500, ecoli.1000`, etc. The second set of datasets consisted of 1, 2, 4, 8, 16, 32, and 46 concatenated assemblies from the Human Pangenome Reference Consortium (HPRC) Year 1 version 2 data freeze (https://human-pangenomics.s3.amazonaws.com/index.html). These datasets will be refered to as HPRC.1, HPRC.2, HPRC.4, etc. All files were provided in FASTA format, which was the file type supported by all tools evaluated. The number of characters, and file sizes for all datasets are in Table 2 and Table 3.

**Table 2.** *E. coli* dataset sizes: total characters and file sizes for different numbers of assemblies.

| # Assemblies | # Characters | Size (GB) |
| --- | --- | --- |
| 500 | 2,548,889,508 | 2.59 |
| 1,000 | 5,128,946,115 | 5.21 |
| 2,000 | 10,291,604,059 | 10.46 |
| 3,682 | 18,942,234,599 | 19.25 |

**Table 3.** Human Pangenome dataset sizes: total characters and file sizes for different numbers of assemblies.

| # Assemblies | # Characters | Size (GB) |
| --- | --- | --- |
| 1 | 9,119,164,921 | 9.27 |
| 2 | 15,086,634,229 | 15.34 |
| 4 | 27,143,994,500 | 27.60 |
| 8 | 51,311,492,262 | 52.17 |
| 16 | 99,654,503,452 | 101.32 |
| 32 | 196,379,510,817 | 199.66 |
| 46 | 277,515,117,856 | 282.15 |

### 4.2. Experimental Set-up

We compared `RAPID-PFP` to `PFP-FL` [15] and `PFP-ITL` [5], two established implementations of the `PFP` algorithm. We briefly note that `PFP-ITL` is part of a larger method called BigBWT that aims to build the BWT from `PFP`. We also do not execute the entire BigBWT method but only the `newscan` executable which computes only the `PFP` of the input. Additionally, `PFP-FL` was run using multi-threading, but it is not clear to what extent the program is parallelized. All experiments were conducted using Snake-make [12] on nodes with 2.5 GHz AMD EPYC 9655p Turin CPUs running Red Hat Enterprise Linux 9.5 and a 24-hour time limit. For `PFP-FL` and `PFP-ITL`, we used 16 CPUs with 200 GB of RAM, while experiments with `RAPID-PFP` were performed with 1 CPU, an NVIDIA B200 GPU, which has 200 GB of VRAM, and 50 GB of RAM. Runtime and memory usage were measured using the `/usr/bin/time` command, and VRAM usage by `RAPID-PFP` was recorded with the Rapids Memory Manager. All tools required the parameters *w* and *p* as defined by `PFP`, which we set to 15 and 100, respectively, for all experiments.

### 4.3. Performance on E. coli Datasets

The wall-clock time for each method for the *E. coli* experiment is shown in Figure 2a and the specific runtimes are given in Table 4. We found that `RAPID-PFP` consistently required less time than both `PFP-FL` and `PFP-ITL` across all dataset sizes. At 500 concatenated *E. coli* assemblies, `RAPID-PFP` completed in 8 seconds compared to 40 seconds for `PFP-FL` and 172 seconds for `PFP-ITL`, representing a 5.2 times runtime reduction relative to `PFP-FL` and a 22.3 times reduction relative to `PFP-ITL`. The performance gap widened as the input size increased. At 3,682 concatenated assemblies, `RAPID-PFP` (23 seconds) achieved a 32.1 times runtime reduction over `PFP-FL` (552 seconds) and a 62.6 times reduction over `PFP-ITL` (1078 seconds). The speedup of `RAPID-PFP` and `PFP-FL` over `PFP-ITL` can be found in Figure 2b.

**Table 4.** Runtime comparison of RAPID-PFP, PFP-FL, and `PFP-ITL` on *E. coli* datasets reported in hh:mm:ss.mmm. Methods are ordered slowest to fastest within each block, and the fastest runtime is bolded. The final column reports the range of speedups achieved by `RAPID-PFP` over the other tools.

| Dataset | Samples | Method | Runtime (hh:mm:ss.mmm) | RAPID-PFP speedup |
| --- | --- | --- | --- | --- |
| E. coli | 500 | PFP-ITL | 00:02:51.680 | **5.2x–22.3x** |
|  |  | PFP-FL | 00:00:40.110 |  |
|  |  | RAPID-PFP | **00:00:07.710** |  |
| E. coli | 1000 | PFP-ITL | 00:06:39.550 | **7.4x–44.1x** |
|  |  | PFP-FL | 00:01:06.630 |  |
|  |  | RAPID-PFP | **00:00:09.060** |  |
| E. coli | 2000 | PFP-ITL | 00:09:22.780 | **20.3x–38.2x** |
|  |  | PFP-FL | 00:04:59.910 |  |
|  |  | RAPID-PFP | **00:00:14.740** |  |
| E. coli | 3682 | PFP-ITL | 00:17:57.650 | **32.1x–62.6x** |
|  |  | PFP-FL | 00:09:11.750 |  |
|  |  | RAPID-PFP | **00:00:17.210** |  |

**Figure 2:**
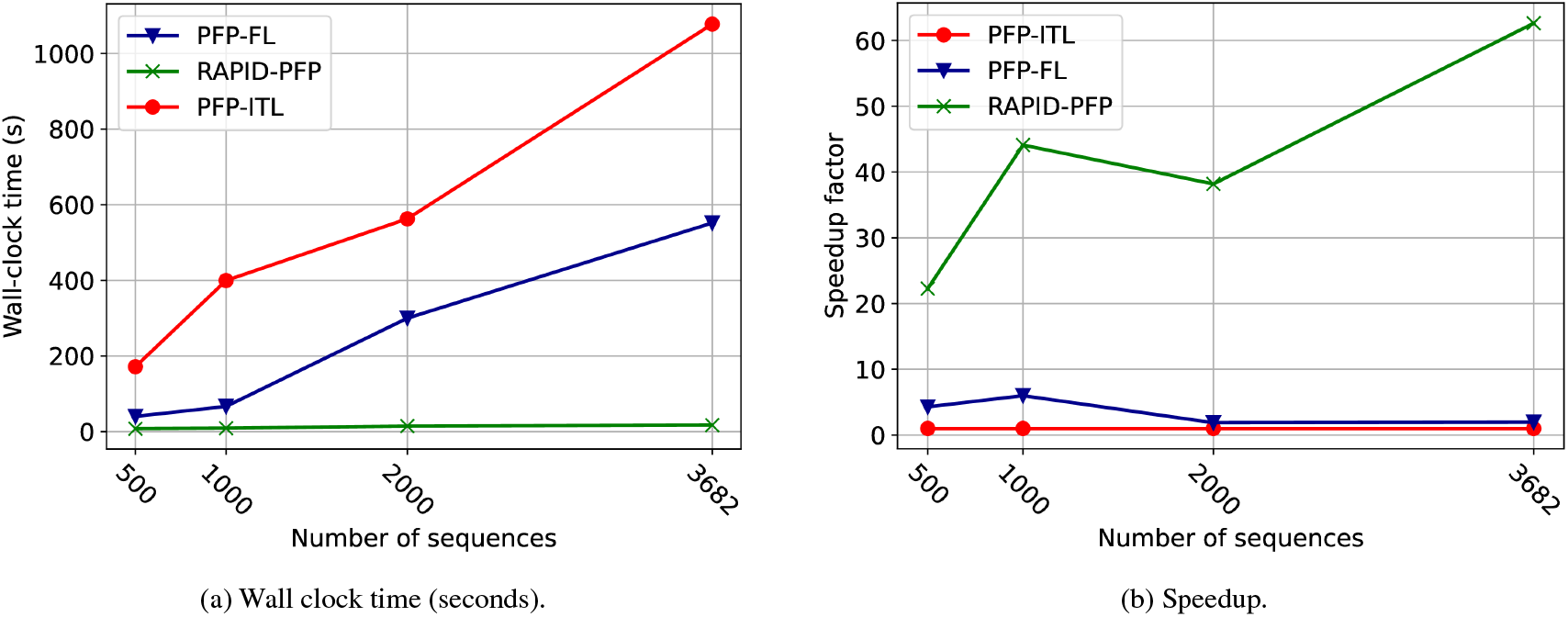
Performance of RAPID-PFP, PFP-FL, and PFP-ITL on *E. coli* datasets from NCBI [2]. Each dataset consists of concatenated assemblies (500, 1000, 2000, and 3,682) in FASTA format. (a) Wall-clock time for Prefix-Free Parsing (PFP) construction. (b) Speedup of RAPID-PFP and PFP-FL relative to the PFP-ITL baseline as a function of the number of samples. Speedup is computed as *T*_PFP-ITL_ *∕T*_*method*_ using wall-clock time.

### 4.4. Performance on Human Pangenome Datasets

We present the wall-clock time plot for the human pangenome experiments in Figure 3a and the speedup plot in Figure 3b. We note that `PFP-ITL` was unable to process datasets larger than four samples due to hash collisions at larger input sizes. As in the experiments with *E. coli*, `RAPID-PFP` consistently outperformed both `PFP-FL` and `PFP-ITL` across all datasets. For a single human genome, `PFP-FL` and `PFP-ITL` performed comparably, requiring 550 and 583 seconds, respectively, while `RAPID-PFP` completed in 40 seconds, which is 13.5 times faster. At four human genome samples, the largest dataset that `PFP-ITL` could process, `RAPID-PFP` required 68 seconds compared to 604 seconds for `PFP-FL` and 2,974 seconds for `PFP-ITL`, achieving a runtime 8.8 times faster relative to `PFP-FL` and 43.1 times faster relative to `PFP-ITL`. In the largest dataset tested (46 human genome samples), `PFP-FL` required 8,655 seconds, while `RAPID-PFP` completed in 359 seconds, yielding a runtime 33.4 times faster. These results emphasize the architectural advantages of `RAPID-PFP`. Unlike CPU-bound implementations, RAPID-PFP exploits parallelism to distribute the prefix-free parsing construction workload across thousands of CUDA cores. The specific runtimes for each method for each sample size is given in Table 5.

**Table 5.** Runtime comparison of RAPID-PFP, PFP-FL, and `PFP-ITL` on `HPRC` datasets reported in hh:mm:ss.mmm. Methods are ordered slowest to fastest within each block, and the fastest runtime is bolded. The final column reports the range of speedups achieved by RAPID-PFP over the other tools.

| Dataset | Samples | Method | Runtime (hh:mm:ss.mmm) | RAPID-PFP speedup |
| --- | --- | --- | --- | --- |
| HPRC | 1 | PFP-ITL | 00:09:42.980 | **13.5x–14.3x** |
|  |  | PFP-FL | 00:09:09.940 |  |
|  |  | RAPID-PFP | **00:00:40.750** |  |
| HPRC | 2 | PFP-ITL | 00:36:11.610 | **15.6x–49.4x** |
|  |  | PFP-FL | 00:11:25.980 |  |
|  |  | RAPID-PFP | **00:00:43.960** |  |
| HPRC | 4 | PFP-ITL | 00:49:33.610 | **8.8x–43.1x** |
|  |  | PFP-FL | 00:10:03.500 |  |
|  |  | RAPID-PFP | **00:01:08.950** |  |
| HPRC | 8 | PFP-FL | 00:33:21.030 | **22.5x** |
|  |  | RAPID-PFP | **00:01:29.110** |  |
|  |  | PFP-ITL | – |  |
| HPRC | 16 | PFP-FL | 00:52:39.820 | **23.6x** |
|  |  | RAPID-PFP | **00:02:14.090** |  |
|  |  | PFP-ITL | – |  |
| HPRC | 32 | PFP-FL | 01:45:12.000 | **30.6x** |
|  |  | RAPID-PFP | **00:03:26.300** |  |
|  |  | PFP-ITL | – |  |
| HPRC | 46 | PFP-FL | 02:24:15.000 | **33.4x** |
|  |  | RAPID-PFP | **00:04:19.110** |  |
|  |  | PFP-ITL | – |  |

**Figure 3:**
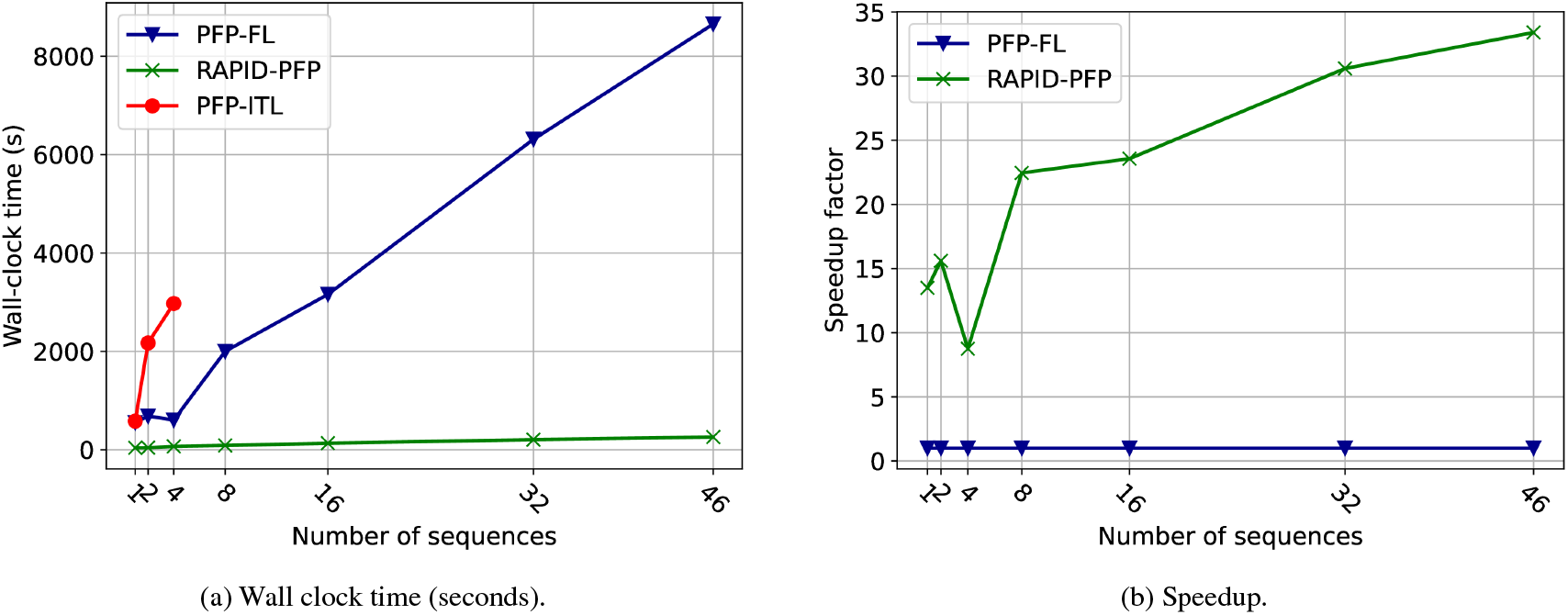
Performance of RAPID-PFP, PFP-FL, and PFP-ITL on human pangenome datasets from the Human Pangenome Reference Consortium (HPRC) Year 1 v2 data freeze (https://human-pangenomics.s3.amazonaws.com/index.html). (a) Wall-clock time for Prefix-Free Parsing (PFP) construction across increasing numbers of concatenated human assemblies (1–46). (b) Speedup of RAPID-PFP relative to the PFP-FL baseline as a function of the number of samples. Speedup is computed as *T*_PFP-FL_ *∕T*_RAPID-PFP_ using wall-clock time. PFP-ITL was omitted from this speedup plot as it failed to complete on all sample sizes.

### 4.5. Detailed Runtime Analysis of RAPID-PF

We briefly present a detailed analysis of the runtime of different steps in `RAPID-PFP` on the HPRC datasets to highlight the efficiency of the GPU operations and identify where bottlenecks lie. A breakdown of the percentages of the total runtime that steps in our algorithm take are shown in Table 6. It is clear from this table that the I/O operations in `RAPID-PFP` constitute a large percentage of the total runtime. For `HPRC.1`, it makes up over 90% of the total runtime and that only decreases to 74% of the total runtime for `HPRC.46`. Part of this I/O, specifically the parquet writing and reading steps, is an inherent aspect of our algorithm which operates on chunks of the input at a time and writes out temporary results to be able to work within the GPU memory. This increases as a percentage of the total runtime as the input size increases, because `RAPID-PFP` needs to work on more input chunks to process the entire input. However, other I/O steps (input reading and dictionary writing), also make up large percentages of the runtime and have the potential to be resolved. For example, in `HPRC.46` the parquet writing step makes up a larger percentage than the dictionary writing step, however, at this file size there are 15 parquet files being written and read in and the dictionary writing step, when writing in the PFP format, takes up just under half as much time with a single write. If instead, the dictionary is written out in a parquet format, it only accounts for 2.7% of the total runtime. Similarly, the input reading step makes up a significantly larger percentage of the total runtime than the temporary parquet I/O step takes. The importance in this is that in the temporary parquet I/O step and when writing the dictionary in parquet format, the I/O is being done in a GPU friendly format, while the other I/O steps are using formats that are not inherently efficient for the GPU. As the use of GPUs in genomic pipelines become more prevalent, the adoption of GPU friendly file formats will allow for `RAPID-PFP` to provide even greater speedups over CPU implementations. When not considering the input reading step, dictionary writing, or parse writing steps, `RAPID-PFP` only takes 153 seconds on HPRC.46, which is an 56 times speedup over `PFP-FL`. This table also highlights the relatively low cost of the custom kernel and GPU operations, which do not scale quickly as the input size increase. This accentuates `RAPID-PFP`’s ability to scale to larger input sizes than would be feasible with current CPU implementations.

**Table 6.** Percentage of total runtime spent in each stage of RAPID-PFP as a function of the number of samples of the HPRC dataset. Values are percentages (%). Each number of samples is shown as a two-row block by output method.

|  | Dictionary<br>Output<br>Format | Input<br>Reading | Trigger<br>String<br>Finding | Expand<br>Sequence | Temp<br>Dictionary<br>Construction | Temp<br>Parquet<br>I/O | Final<br>Dictionary<br>Construction | Dictionary<br>Writing | Parse<br>Construction | Parse<br>Writing |
| --- | --- | --- | --- | --- | --- | --- | --- | --- | --- | --- |
| 1 | PFP | 4.70 | 0.65 | 1.81 | 0.00 | 0.00 | 3.14 | 88.61 | 0.46 | 0.63 |
|  | Parquet | 19.36 | 2.67 | 7.47 | 0.00 | 0.00 | 12.96 | 53.04 | 1.91 | 2.59 |
| 2 | PFP | 7.18 | 0.58 | 2.62 | 0.00 | 0.00 | 4.27 | 83.01 | 1.50 | 0.84 |
|  | Parquet | 22.64 | 1.83 | 8.26 | 0.00 | 0.00 | 13.49 | 46.37 | 4.75 | 2.66 |
| 4 | PFP | 7.23 | 0.41 | 2.72 | 3.68 | 18.62 | 1.28 | 52.68 | 12.47 | 0.91 |
|  | Parquet | 13.25 | 0.76 | 4.98 | 6.75 | 34.12 | 2.34 | 13.29 | 22.85 | 1.66 |
| 8 | PFP | 9.93 | 0.44 | 3.80 | 5.23 | 22.67 | 1.67 | 41.49 | 13.46 | 1.31 |
|  | Parquet | 15.41 | 0.68 | 5.90 | 8.11 | 35.16 | 2.60 | 9.22 | 20.88 | 2.04 |
| 16 | PFP | 12.82 | 0.77 | 4.97 | 7.01 | 28.15 | 1.92 | 28.96 | 13.84 | 1.56 |
|  | Parquet | 17.04 | 1.03 | 6.60 | 9.32 | 37.42 | 2.54 | 5.57 | 18.40 | 2.08 |
| 32 | PFP | 16.05 | 0.70 | 6.09 | 8.60 | 33.48 | 2.29 | 19.33 | 11.49 | 1.97 |
|  | Parquet | 19.17 | 0.83 | 7.27 | 10.27 | 40.00 | 2.74 | 3.63 | 13.73 | 2.36 |
| 46 | PFP | 18.13 | 0.72 | 6.78 | 9.67 | 37.48 | 2.93 | 15.67 | 5.85 | 2.77 |
|  | Parquet | 20.90 | 0.83 | 7.81 | 11.16 | 43.21 | 3.38 | 2.77 | 6.75 | 3.19 |

## 5. Conclusion

In this paper, we introduced `RAPID-PFP`, a GPU-accelerated implementation of Prefix-Free Parsing (PFP) designed to meet the computational demands of large-scale repetitive datasets. Using the parallelism of modern GPU architectures and optimizing critical components such as parse construction and dictionary building, `RAPID-PFP` achieves substantial performance gains over existing CPU-based implementations. Our experimental results demonstrate up to 45 times speedup compared to `PFP-FL` and `PFP-ITL`, enabling the construction of PFPs on datasets of unprecedented size, including thousands of concatenated *E. coli* assemblies and the complete Human Pangenome Reference Consortium (HPRC) dataset. These results show the potential of GPU acceleration to address the growing computational challenges of pangenomics and other high-throughput applications.

The design of `RAPID-PFP` reflects several key engineering insights. First, the batching strategy and memory management effectively mitigate GPU memory constraints, supporting efficient processing of large inputs even on a single device. Second, the use of custom CUDA kernels is used to parallelize fine-grained steps of a previously sequential algorithm. This highlights the value of rethinking the foundational algorithms for parallel hardware.

Looking ahead, several directions remain to further enhance the scalability and usability of `RAPID-PFP`. One avenue is the integration of multi-GPU support, which would distribute memory and computation across multiple devices, enabling the processing of even larger pangenomic datasets and further reducing wall-clock time. Another promising direction is the development of GPU-accelerated pipelines for downstream applications of PFP, such as Burrows-Wheeler Transform (BWT) construction, suffix array generation, and FM-index building. Finally, the development of GPU friendly genomic file types would significantly reduce the amount of time necessary for I/O operations in `RAPID-PFP`, allowing for even greater speedups. These extensions would enable end-to-end GPU-based workflows for constructing succinct data structures on massive genomic collections.

## 6. Acknowledgments

This work is supported in part by funds from the National Science Foundation (NSF: # 1636933 and # 1920920).

## CRediT authorship contribution statement

**Eddie Ferro:** Conceptualization, Data Curation, Validation, Software, Writing - original draft, Methodology, Formal Analysis. **Tyler Pencinger:** Software. **Oded Green:** Writing - review & editing, Resources. **Mahsa Lotfollahi:** Resources. **Christina Boucher:** Conceptualization, Supervision, Methodology, Project Administration, Writing - review & editing, Resources.

## 7. Data Availability

All of our code is publicly available on GitHub at https://github.com/EddieFerro/gpuPFP. Data will be made available upon request.

